# Transparent and Conformal Microcoil Arrays for Spatially Selective Neuronal Activation

**DOI:** 10.1101/2021.12.07.471184

**Authors:** Vineeth Raghuram, Aditya D. Datye, Shelley I. Fried, Brian P. Timko

**Affiliations:** Dept. of Biomedical Engineering, Tufts University, Medford, MA 02155, USA; Boston VA Healthcare System, Boston, MA 02130, USA; Dept. of Neurosurgery, Massachusetts General Hospital, Harvard Medical School, Boston, MA, 02114, USA

## Abstract

Micromagnetic stimulation (μMS) using small, implantable microcoils is a promising modality for achieving neuronal activation with high spatial resolution and low toxicity. Microcoils can be designed to achieve localized, spatially asymmetric fields that target neurons of a particular orientation. Insulation of the coil avoids the direct contact between metal and tissue and the use of specialized biopolymers may help to further reduce chronic inflammation and glial scarring. Studies to date have largely focused on single channel devices; herein, we report the design and development of a microcoil array for localized activation of cortical neurons and retinal ganglion cells. We utilized a computational model that related the activation function to the geometry and arrangement of coils and selected a coil design that maintained a region of activation <50 µm wide. The device was composed of an SU8/Cu/SU8 tri-layer structure, which was flexible, transparent and conformal and featured four individually-addressable microcoil stimulation elements. Interfaced with ex vivo cortex or retina slices from GCaMP6-transfected mice, we observed that individual neurons localized within 40 µm of the element tip could be activated repeatedly and in a dose (power) dependent fashion. Taken together, these results highlight the potential of magnetic stimulation devices for brain-machine interfaces and could open new routes toward bioelectronic therapies including prosthetic vision devices.

## Introduction

Implantable devices that achieve localized neuronal activation hold great potential in bioelectronic medicine, with applications such as treating neurological or mood disorders as well as restoring vision (1) or other sensory deficits (2). Stimulation is typically achieved with multi- electrode arrays (MEAs), including Michigan- and Utah-style arrays which are fabricated on rigid substrates and have been demonstrated intracortically. Compared to noninvasive stimulation techniques such as Transcutaneous Electrical Nerve Stimulation (TENS), these implantable devices can better confine activation, e.g., tens to hundreds of microns around the electrode site for implants vs. hundreds of microns or millimeters for noninvasive. More recently, MEAs have been demonstrated on soft, stretchable and conformal substrates (3, 4). Such an approach enables all electrodes within the array to be in close proximity to complex 3-d neuronal structures, e.g., the innermost surface of the retina (5); or embedded within engineered tissues which may serve a role in disease modeling and regenerative medicine (6, 7).

Conventional stimulation devices are composed of metallic electrodes that activate neurons through capacitive currents. While widely explored, this paradigm has several key drawbacks. First, the metallic electrodes present a large mechanical mismatch with brain tissue (Young’s modulus E>10 GPa for gold vs. 1 kPa for brain). This mismatch causes inflammation and oxidative stress at chronic time points (8), as well as glial scarring which ultimately limits the lifetime of the device. Second, prolonged currents themselves can also cause neuronal damage, with some stimulation regimes causing significant loss within 60 μm of the electrode after 30 days stimulation (9). Finally, these devices generate fields whose size and shape cannot be readily controlled, especially in inhomogeneous or anisotropic tissues which cause irregular distribution of the electric field (10). While some studies have reported differences in activation thresholds that can be exploited to selectively activate certain neuronal sub-populations (11–13), tissue- independent control over the shape of the field to activate neurons in a certain orientation would enable further selectivity. Taken together, these limitations present significant roadblocks toward the widespread adoption of commercial implants (14, 15).

Microcoils that magnetically activate neurons could offer significant advantages over conventional metallic devices. Since magnetic fields readily permeate both tissues and biomaterials, microcoils could provide for much more reproducible activation in inhomogeneous tissues (16, 17). They could also be completely encapsulated in one or more biopolymer layers to avoid direct neuron/metal interactions. These coatings could include mechanically-distinct interlayers, which were recently used to achieve bioelectronics with tissue-level moduli and minimal foreign-body responses (18), or a bioactive layer to further enhance tissue integration (19). Chronic magnetic stimulation has also been shown to be minimally toxic to neurons (10). Furthermore, magnetic coils generate localized, spatially asymmetric electric fields whose shape can be tuned through the geometry of the microcoil. These asymmetric fields are thought to enable selective targeting of specific neuronal sub-populations, for example vertically oriented pyramidal neurons while avoiding horizontally aligned passing axon fibers (20). Such functionality allows for more focal confinement of activation and thus could allow for the restoration of high acuity vision or enable activation in specific regions of the somatosensory cortex, e.g., to enhance feedback as part of a brain-machine interface system.

Prior studies have reported single microcoil devices that could generate small, focal regions of activation (21) and have explored the effects of varying coil shape and design, on the both the spatial selectivity and strength of activation (22). While devices that consist of a single microcoil may be useful in targeting one isolated region of the cortex, the ability to focally activate multiple cortical columns, or specific cortical layers, either in synchrony or in temporally modulated intervals (interleaved), would improve the translational value for clinical applications where the creation of multiple regions of neural activity is the goal.

In this study, we report a flexible, transparent, and conformal bioelectronic device that includes an array of four micromagnetic stimulation (μMS) elements. The components are composed of copper coils and encapsulated within SU-8, a photo-crosslinkable, bioinert material that has been demonstrated in chronic bioelectronic recording studies (23, 24). We develop a computational model to explore the relationship between the shape of the microcoil and resulting activating function. We then demonstrate localized activation in ex vivo tissue slices from the retina and cortex, both of which are associated with vision and are active areas of study for prosthetic devices that address blindness. Our technology represents a general platform that could be scaled to multiplexed arrays, broadened to incorporate other bioelectronics such as recording elements (23, 24), and extended to other substrate geometries to enable, for example, injectable and minimally-invasive devices (25–27).

## Results

### Computational model for coil design

We started by developing a computational model (28) (see Methods) to explore the interaction between the electric fields induced by two adjacent coils (Figure 1). Such interactions have not been well studied and so we set out to (i) explore the relationship between coil spacing and field strength with the goal of determining an appropriate inter-coil spacing for the new array, and (ii) test the influence of coil shape on the interactions between neighboring coils. Figure 1A shows the 3 tip designs evaluated here: rectangular, U-shaped, and V-shaped. Similar to previous studies (21, 22, 29), the region surrounding the arrays was modeled as a homogenous, isotropic medium with the properties of grey matter. In order to isolate the effect of coil geometry, the distance between the vertical leads of the coils in each array was fixed at 100 μm and the rate of change of current through the coils was maintained at 1 A/μs (direction of current in each coil is indicated by arrows).

**Figure 1.**
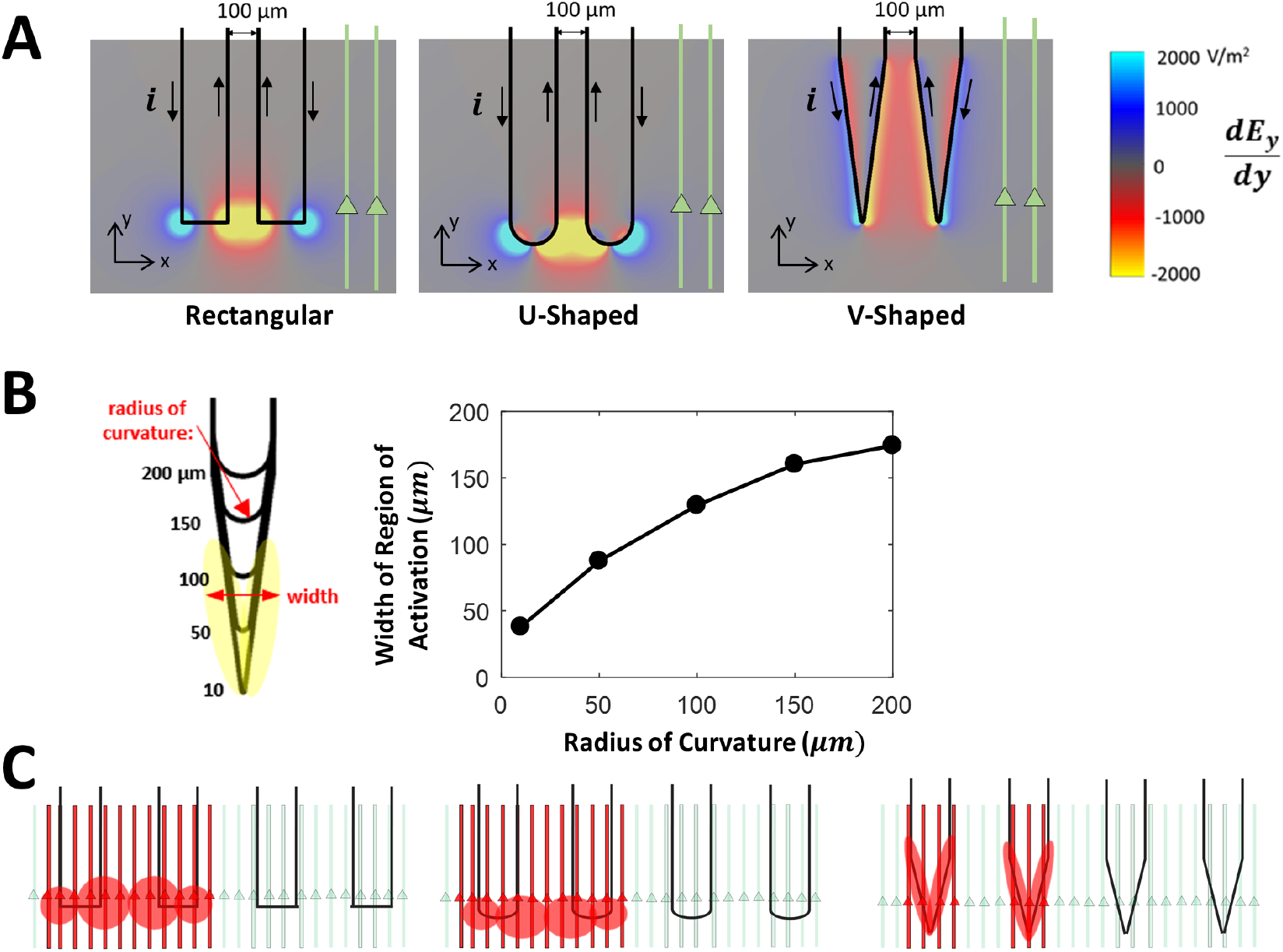
Computational modeling of microcoil array geometries and experimental schematic (A) Heat maps depicting the induced electric field gradient produced by rectangular, U- and V- shaped geometries. Gradients are calculated on a plane that is 10 μm below the bottom edge of the array and with di/dt=1A/μs. Black arrows depict the directions of current through the coils. (left) Schematic illustration of a microcoil with a gradually increasing radius of curvature at its tip. (right) Relationship between bend radius of the tip and width of the region of activation field strength > 2000 V/m^2^. (C) Schematic illustration of pyramidal neuron axons (light green) interfaced with devices and corresponding activation regions (red; adapted from the yellow regions shown in panel a). Axons passing through regions of stronger gradient are more likely to be activated (red), therefore, the model predicts that V-shaped devices will offer better spatial resolution.

Previous computational studies (30–32) have shown that the strength of the gradient of the electric field arising along the length of a targeted axon (referred to as the activating function (30)) is a good predictor of the effectiveness of a given set of μMS conditions. Because the axons of cortical pyramidal neurons are generally oriented in the same direction (two sample cells/axons are depicted), we started by determining the spatial gradient of the induced electric field along the length of these axons 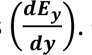. Gradients were calculated along a plane 10 μm below the surface of the array and are shown in the colormaps of Figure 1A. The maximum gradient levels in the maps were fixed at ±2000 V/m^2^ in order to facilitate comparisons across the three coil geometries; saturated yellow and blue regions correspond to negative (< -2000 V/m^2^) and positive (> +2000 V/m^2^) gradients (respectively).

When the coil shape was rectangular (Figure 1A, left), gradients were strongest at the two right-angle bends and decreased radially with distance from the corner, thereby resulting in circular- shaped regions. The polarity of the gradient was dependent on the direction of flow of the current in the vertical portions of the microcoil and thus one gradient was positive and the other negative.

Narrowing the width of the coil (the space between its ascending and descending vertical portions brought the opposing field gradients close to each other and resulted in reductions of the peak magnitude for each (not shown). However, because the electrical current flowed in opposite directions for the 2 adjacent coils (black vertical arrows), the polarity of the induced field gradients were the same in the region between coils and thus there was constructive interference, and the combined field was broader and stronger. This is the same principle used for figure-8 coils in transcranial magnetic stimulation (TMS). The width of the region for which gradient exceeded 2000 V/m^2^ was larger than the 100-μm separation between coils. A qualitatively similar region of activation was produced when the bends in the coil tips were U- shaped (Figure 1A, middle), along with a similar overlap in the central region that both strengthened and broadened it. When the coils were V-shaped (Figure 1A, right), the regions of peak gradient were narrower, in part because the close proximity of the ascending and descending branches at the tip resulted in overlap of the positive and negative regions of field gradient. As a result, the regions of peak gradient were narrowly confined to the tip region of each coil and there was little overlap. Thus, even though the sharper bend associated with V-shaped coils produces a stronger local field, the increased separation between adjacent tips reduces the potential for overlap. Taken together, these results suggest the use of V-shaped coils may be more effective in creating multiple, distinct regions of neural activation. In contrast, the expansive regions and overlap produced by the rectangular and U-shaped designs might result in reduced channel independence.

The relationship between coil geometry and the resulting region of strong field gradient was further explored by systematically varying the radius of curvature of the coil bend at the tip (this resulted in a gradual transition from a V- to U-shaped design) (Figure. 1B, right): we again calculated the width of the region of activation where *d*Ey/*d*y < -2000 V/m^2^. Consistent with the results of Figure 1A, a smaller bend radius was associated with a narrower region of peak gradient. This pattern is also consistent with the analogous relationship between the radii of coils utilized in TMS and their induced fields: TMS coils with smaller dimensions generally produce electric fields with a smaller tangential spread (33). Because our goal was to minimize the overlap between the adjacent coils, while maintaining channel independence, we chose to proceed with V-shaped coil tips for subsequent physiological experiments.

### Fabrication, assembly and testing of microcoil arrays

We designed and fabricated an array with four adjacent microcoils, with the tips equally spaced 250 µm apart (Figure 2A,B). This distance was chosen as it approximates the distance between adjacent cortical columns (34). The device consisted of Cu coils and interconnects encapsulated by two layers of SU-8, which functioned both as flexible support substrate and passivation layer. A series of eight copper pads, with each set of two adjacent pads corresponding to the signal and ground elements for an individual coil, allowed the four coils to be individually addressed. A distance of 3.5 cm between the rectangular layers of interconnects and the microcoils allowed for facile placement of the functional portion of the coil onto the microscope stage as well as sufficient distance between the base of the device and the chamber slide on which the tissue slice would be placed. The dimensions of the multi-coil array were specifically chosen for use in *in-vitro* electrophysiology experiments that would be used to assess functionality of the design. Given that the width of a typical adult mouse in-vitro cortical slice is around 4.5 mm and the total length between the edges of the substrate is about 1.3 mm, the multicoil array would occupy only about 30% of the tissue width and could be easily rotated to adapt to different slice orientations. As a typical pyramidal neuron has a soma diameter of around 20-25 µm, the distance between the coils should allow the visualization of activation up to 10 or 11 cells. To evaluate device transparency, we interfaced the array on-top of cortical tissue in both fluorescence and brightfield mode (Figure 2C) and found clear visualization of the tissue underneath the device. This provided confirmation that our device would be suitable for simultaneous calcium imaging applications.

**Figure 2.**
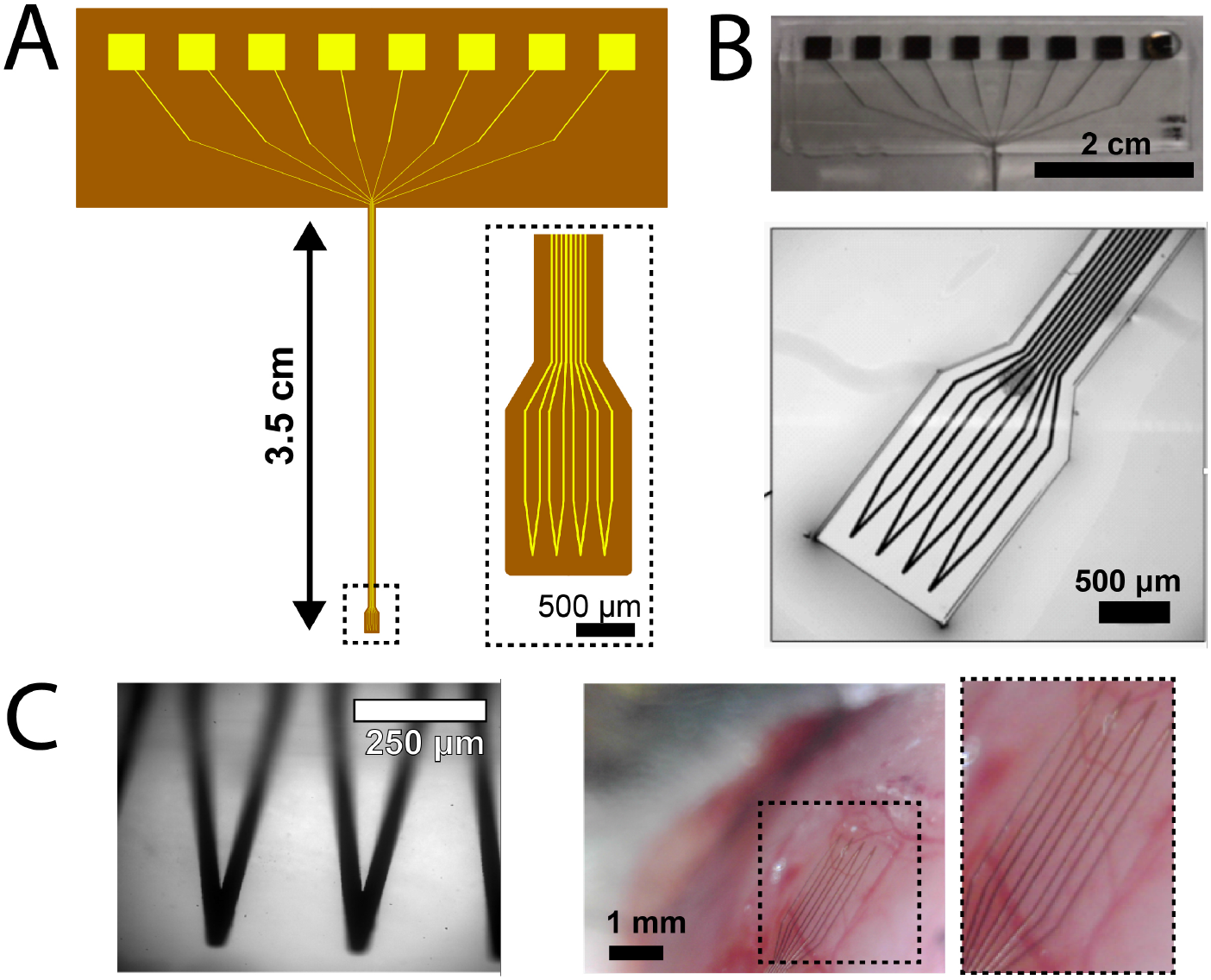
Fabrication and assembly of flexible microcoil devices (A) Schematic of multicoil array. Each pair of square pads addresses 1 coil. Inset, expanded region of stimulation region with 4 microcoils. (B) Photographs of (top) interconnect and (bottom) simulation regions. (C) Microcoil array interfaced with cortical tissue in (left) fluorescence and (right) brightfield mode. Inset, expansion of device and underlying tissue.

To ensure the fabricated coils would achieve μMS as intended, i.e., generally consistent with the model predictions, we first characterized the electrical integrity of each coil. Coils were submerged into a bath containing PBS solution and the integrity of the SU-8 insulating layer was assessed by measuring its impedance (DC impedance between a lead connected to the coil trace and a lead inserted in the bath but adjacent to the array). Impedance levels were typically above 200 MΩ and often exceed 1 GΩ suggesting good integrity of the SU-8 layer and therefore minimizing the possibility for direct (electrical) activation of neurons by the coil. Next, the integrity of the copper trace was evaluated by measuring the DC impedance across its input and output leads. Measurements were typically <40 Ω suggesting good integrity of the copper trace.

### Validation of microcoils for μMS

Stimulus waveforms were generated by connecting the output of a function generator to an audio amplifier (PB 717X, 1000W, Pyramid Inc.) with a gain of 5.6V/V and a bandwidth of 70 kHz (21). The function generator was programmed to deliver a continuous series of ramp pulses with amplitudes ranging from 0 to 1500mV; the duration of both the rising and falling phases was 0.1ms (total 0.2 ms) and there was a 5 ms interval between the offset of one ramp waveform and the onset of the next one. A glass patch pipette was positioned 5 µm above the plane of one of the fabricated coils to measure the response in the bath resulting from the flow of current through the coil (Figure 3A,B). The signal captured by the patch pipette (Figure 3C) is a measure of the current injected by the patch amplifier to offset the change in voltage at the pipette tip arising from the flow of current through the microcoil (see Methods). Responses typically consisted of a series of biphasic waveform with positive and negative phases corresponded to the rise and falling elements of the input ramp pulse (Figure 3C, right), consistent with similar types of measurements made previously with both bent wire and fabricated coils (20). As expected, the amplitude of the response increased linearly with the amplitude of the stimulus current (Figure 3D). Note that the amplitude of the stimulus currents referred to above, as well as the rest of the manuscript, refer to the output of the audio amplifier.

**Figure 3.**
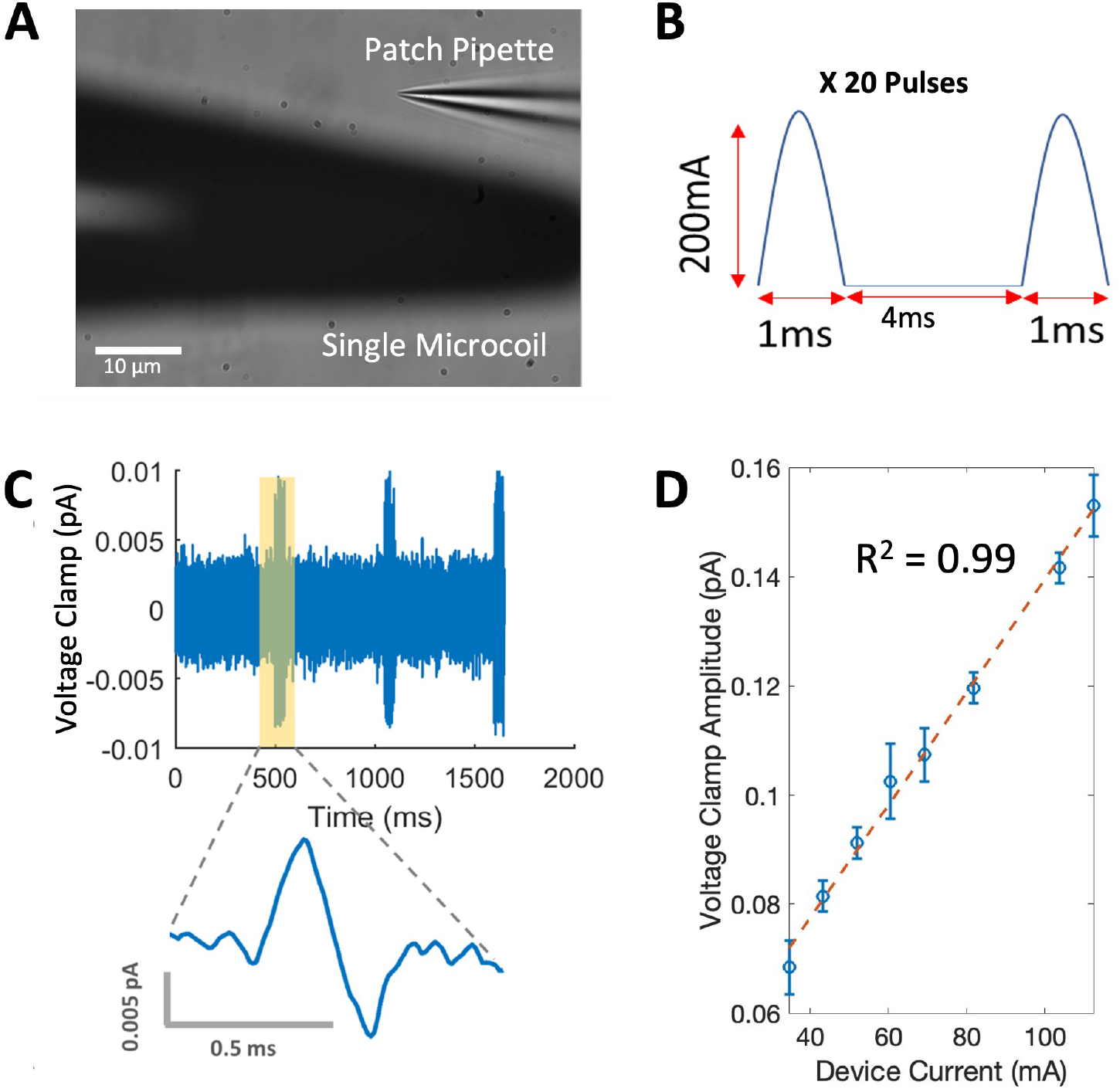
Electrical validation of flexible microcoil devices (A) Photograph of patch clamp pipette positioned 10 µm above one of the V-shaped microcoils in the fabricated array. (B) Schematic of the half-sinusoidal waveform used to test electrical integrity (C) (top) Raw recording of the recorded response to 3 trains of 20 half-sinusoids. (bottom) Expanded view of a typical single (biphasic) response waveform. (D) Plot of the amplitude of the response waveform as a function of the amplitude of the stimulus current (n=3). Error bars represent S.D.

### Flexible microcoil arrays effectively activate CNS neurons

To evaluate the efficacy of μMS with the new coil array, we performed a series of physiological experiments using brain slices obtained from the visual cortex of Thy-1 GCaMP6f mice (20–22). In these animals, a genetically encoded fluorescent calcium indicator labels several neuronal populations within the CNS (including pyramidal neurons in the cortex). The fluorescence level changes systematically with the level of neuronal activation and thus facilitates the study of responses from large numbers of neurons simultaneously (20–22). The microcoil array was overlaid on the surface of the brain slice (ca. 300 µm thickness) so that the response of layer 5 pyramids neurons to magnetic stimulation could be visualized (Figure 4A). To observe activation of individual pyramidal neurons, we delivered a series of stimuli, each consisting of 20 half-sinusoidal waveforms (Fig. 4Ai; the 1ms duration of each corresponded to a sinusoidal frequency of 500 Hz) with an interval of 4ms between two consecutive stimuli (corresponding to a delivery rate of 200 waveforms per second). 9 ‘bursts’ of 20 half sinusoids were delivered in a single trial with 500 ms between consecutive bursts (Figure 4Ai). These parameters were similar to those used in previous in vitro and in vivo microcoil studies (22, 35). The amplitude of individual waveforms was varied to establish the minimum level at which a calcium response was produced within individual cell bodies (identified off-line during post processing). We utilized the NIH ImageJ/FIJI software package to quantify the change in fluorescence that was observed individually within each neuron. We found that each individual stimuli typically elicited a neuronal calcium response (Figure 4Aii,iii), thereby establishing the ability of a single microcoil in our array to activate neurons.

**Figure 4.**
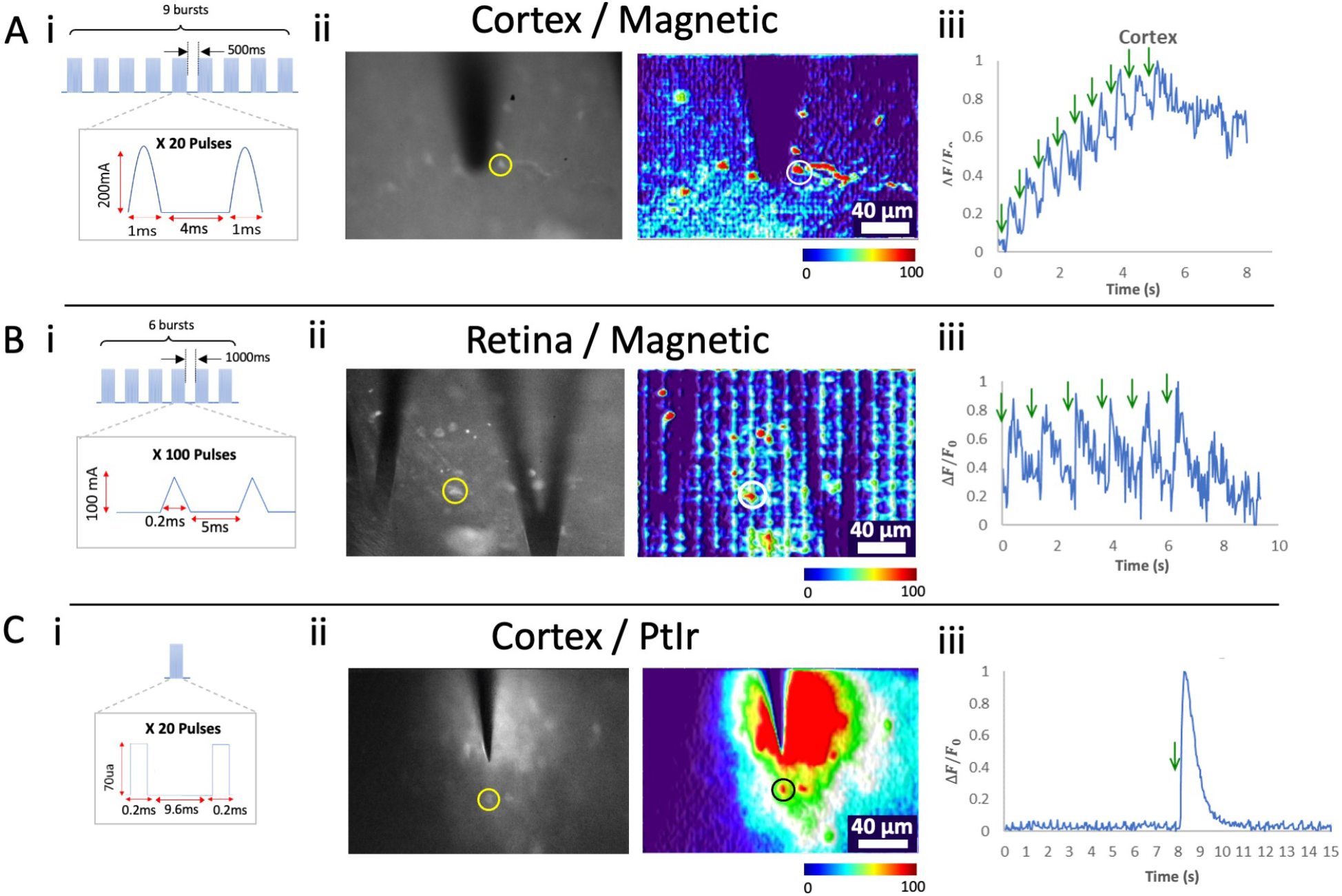
Stimulation and calcium fluorescence recording with *ex vivo* cortex and retina slices. (A) Cortex slice interfaced with microcoil stimulation device. (B) Retina slice interfaced with microcoil stimulation device. (C) Cortex slice interfaced with 10-kΩ PtIr microelectrode. Sub- panels show: (i) Schematic of stimulus pulse train. (ii) (left) fluorescence image and (right) heat map of stimulated tissues. Both images represent peak fluorescence. (iii) Plot of normalized fluorescence vs. time corresponding to the individual neurons indicated by circles in ii. Green arrows indicate the onset of each waveform burst.

We also evaluated μMS with retinal ganglion cells (RGCs) (Methods). The microcoil array was placed on the top surface of a retinal explant, oriented so that the RGCs were directly underneath the coil traces (Figure 4B). Here, a series of 100 waveforms was delivered through a single coil. Repetitive instances of the 100-waveform train were separated by 1s (Figure 4Bi). For the RGC experiments, the shape of the stimulus waveform was switched from sinusoidal to ramp (triangular) to match previous work in the retina. Similar to the results with cortical neurons, each of the ramp stimuli reliably elicited a calcium response in individual RGCs (amplitude 100 mA, Figure 4Bii,iii).

We compared the responses generated by these microcoils to those generated in response to electric stimulation from microelectrodes, i.e., our previous experiments in the cortical slice with microcoils were repeated using a single 10 kΩ Pt-Ir microelectrode (Figure 4C). Stimulus trains consisted of 20 square pulses (0.2 ms duration) with an interpulse interval of 9.6 ms. There was a 500 ms interval between consecutive trains (Figure 4Ci). The pulses were biphasic and rectangular in shape, each at an amplitude of 70 µA. Stimulus parameters (pulse frequency, shape, interpulse interval) were again chosen based on previous studies (21, 22) while amplitude was increased as needed to elicit responses. The neuronal patterns of responses to μMS were significantly different from microcoils and included greater activation of surrounding neuropil and somata (Figure 4Cii,iii).

To further evaluate the efficacy of our microcoils, we studied the relationship between the amplitude of μMS stimulation waveform and the fluorescence change of cortical neurons (**Fig. 5A**). Dose-response curves are shown for 11 cells - the shaded error bars indicate the standard deviation across repeated trials (n = 3). Following the approach utilized by a previous study (36), we defined threshold as the stimulus amplitude required to elicit 50% of the maximum fluorescence change threshold (*I*_50_) (Methods). The threshold for each cell was then plotted as a function of distance from the tip of the coil (**Fig. 5B**). Somewhat surprisingly, there was only a mild correlation between the distance from the coil and threshold (correlation coefficient: *r* = 0.22). However, given the small number of cells tested and potential confounding factors such as cell type, morphology (11–13), and the relatively small distances evaluated here, further experimentation is needed to better assess this relationship. Interestingly, the change in fluorescence was confined to a small region near the tip of the coil with the maximum distance at which a fluorescence change could be observed was 40 μm.

**Figure 5:**
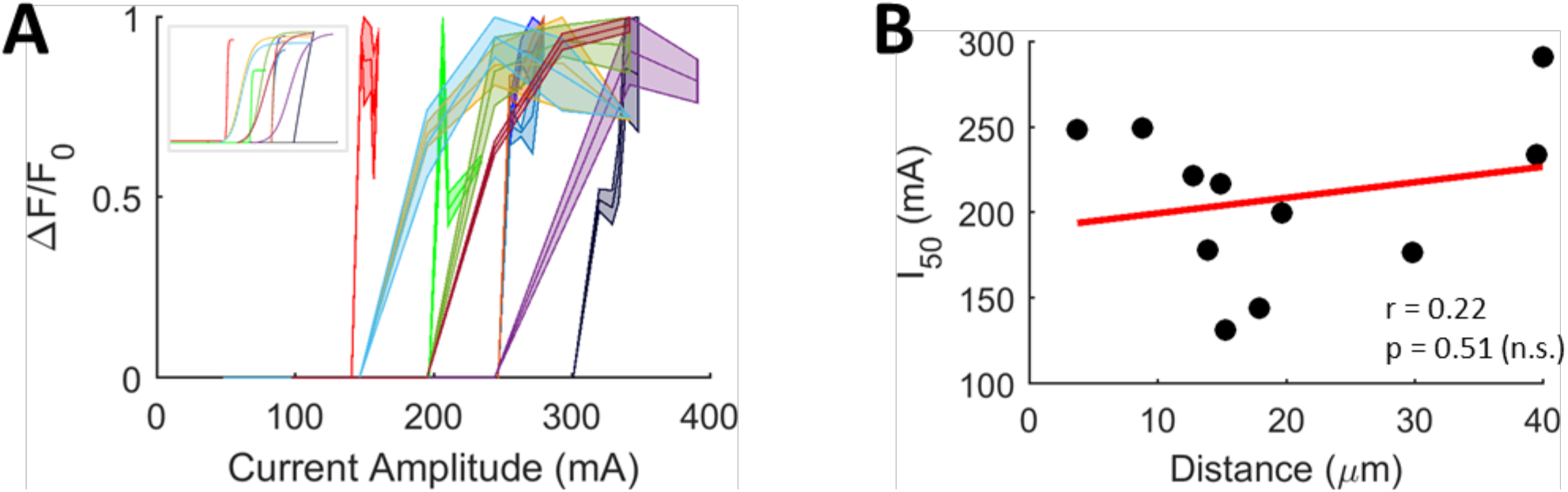
Cortical neurons are sensitive to stimulus amplitude. (A) Dose-response curves depict the relationship between amplitude of magnetic stimulation and (normalized) change in fluorescence of cortical neurons. Each color corresponds to a different cell (11 in total, 5 slices). Shaded error bars indicate the standard deviation across repeated trials (n = 3). *Inset:* Sigmoid fits for each dose-response curve. (B) Relationship between the distance from soma to coil tip and magnetic stimulation threshold (estimated as the current required to elicit 50% of the maximum fluorescence change). Line of best fit (red) and regression analysis are shown.

## Discussion

We designed and fabricated a flexible, transparent, and conformal microcoil device array that could activate neurons by μMS. We showed that gradient of the electric field (activating function) was a monotonic function of the coil radius of curvature; thus by using a coil with a sharp “V” shape we could selectively activate neurons localized within 40 μm of the tip. This resolution represents a significant advance over conventional capacitively-coupled electrodes and could be used to selectively activate individual neuronal pathways.

While the studies in this work were confined to μMS via a single device, the technology could be readily scaled up to hundreds or thousands of channels, as has been achieved with MEAs (37). Aside from achieving highly-multiplexed, high-resolution inputs, μMS is advantageous in that the shape of the activating function can be tuned. We showed that our discrete devices generated anisotropic fields that might be used to selectively activate aligned axons. Alternatively, devices that generate wider activating functions, when multiplexed, are likely to generate interference patterns as we modeled for U- and square-shaped electrodes in Figure 1A. These patterns might be adjusted dynamically, by modulating the amplitude and phase of current passed through each device. Magnetic interference has been used to pattern continuous, complex geometries into composite polymer systems (38); similar techniques might be applied to μMS to encode information at smaller pitch than the electrodes themselves, to further increase the resolution of the technique.

Achieving stable interfaces between devices and tissues has been a major challenge in the bioelectronics field. Our demonstration that SU-8-encapsulated devices can achieve μMS opens new avenues high-resolution bioelectronic stimulation with minimal chronic inflammation. For example, SU-8 meshes injected into living rat brains achieved bioelectronic recordings for at least eight months (39), without disrupting the surrounding neural networks or causing inflammation. Similar meshes have also been wrapped around wrapped around (40, 41) or embedded within (42) the brain organoids in vitro, and in the latter case provided stable bioelectronic readouts for up to 10 months. Very recently, bioelectronic meshes were loaded onto an endovascular catheter, injected into a rat model, and implanted into the middle cerebral artery and anterior cerebral artery, which overlay the cortex and olfactory bulb, respectively. These devices stably recorded neural signals across the vasculature wall, without the need for brain surgery (25, 26). Minimally- or non-invasive bioelectronics could in the future be enabled with μMS, potentially achieving closed-loop systems for neuromodulation.

## Methods

### Microfabrication Protocol

All photoresists and developers were used as received from Kayaku Advanced Materials, Westborough, MA. (i) A 50 nm sacrificial layer of germanium (Ge) was sputtered on onto silicon/silicon oxide substrates (300nm, Cat. 2525). (ii) The bottom passivation layer of the device was composed of 10 μm thick SU8 (SU8 2010) that was spin coated, patterned with UV photolithography (16s @ 15mW/cm^2^) and developed with SU-8 developer for 1 minute. (iii) The metal layer was defined using a LOR20B/S1813 bi-layer stack followed by UV exposure (12s @ 15mW/cm^2^), developed using CD-26, metallized with a 1 μm layer of copper deposited by thermal evaporation (LC Technology Solutions, Salisbury MA), and lifted off with Remover 1165 at room temperature. (iv) The top passivation layer of the device was composed of 20 μm thick SU8 (SU8 2025) that was spin coated and patterned with UV photolithography (21s @ 15mW/cm^2^) with a develop time of 5 minutes. (v) Devices were lifted off by dissolving the Ge layer in DI water at 80°C for up to 48 hours. Next, the interconnect section (Figure 2B, top right) was adhered onto a rigid plastic backing and wires were attached to the exposed copper pads using silver epoxy paste (CW2400, Chemtronics) (Figure 2B,left). The resistance of each device including interconnects was 40-60 Ω and was consistent between consecutive trials.

### Modeling

We developed a computational model to characterize the induced electric fields arising from the flow of electrical current in an array of microcoils consisting of different coil geometries (rectangular, U-shaped, and V-shaped). We utilized a charge-based boundary element fast multipole method (BEM-FMM) toolkit to calculate the induced electric fields within the tissue. Unlike the commonly-applied finite element method (FEM), BEM-FMM operates directly on tissue conductivity boundary surfaces, thus eliminating the need for volumetric meshing (28). After the solution at the tissue boundaries is determined through an iterative process, the magnetic and electric fields from the flow of current through an insulated coil can be evaluated in 3D space, similar to general-purpose FEM software. An important advantage of the BEM-FMM approach is that it allows unconstrained numerical field resolution near tissue interfaces, and thus the induced fields and their spatial gradients can be calculated at a high resolution without the secondary interpolations required for volumetric FEM-based solvers. The BEM-FMM approach is also more computationally efficient compared to existing software packages (28). The region surrounding the array (600 × 600 × 600 μm cube) was modeled as a homogenous, isotropic medium with the properties of grey matter (electrical conductivity: σ = 0.276 S/m) (43), similar to the approach used earlier by members of our group to characterize the spatial extent of intracortical microcoil-based stimulation (22). After the modeling setup was defined, the BEM-FMM toolkit was utilized to calculate the electric field inside the tissue (with a spatial resolution of 1 μm) in response to an applied AC current.

In the quasistatic limit, which is commonly used when modeling electromagnetic fields in biological tissues and is considered valid for frequencies up to ∼10 kHz (44), the total induced electric field is given by:

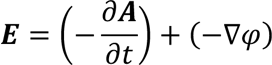

where ***A*** is the magnetic vector potential and φ represents the electric potential due to surface charge redistribution at tissue conductivity boundaries (45). The secondary field (−∇φ) can be ignored if the primary field 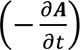 is parallel to the air-tissue boundary (46) as we have done here. The primary field was obtained by discretizing the coil into small, straight elements of current *i*_*j*_(*t*) with orientation ***s***_*j*_) and center coordinate ***p***_*j*_. The magnetic vector potential created by a current element at an observation point ***c***_*i*_ is given by:

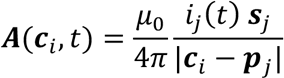

where μ_0_ is the magnetic permeability of vacuum (28). The corresponding electric field generated by the current element is:

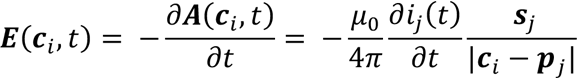

The contributions from each element *j* of the coil are summed to find the total electric field distribution. All calculations were performed at an instant in time when the rate of change of current (di/dt) through the coil was 1A/µs which is a representative value that has been utilized in prior computational studies involving magnetic stimulation (47). Simulations were performed on an Intel Core i7-6700 CPU (3.40 GHz, 32 GB RAM, 4 cores) with an average computation time of ∼1 minute for each simulated microcoil array.

### In-vitro brain slice experiments

Electrophysiological recordings were performed using brain slices prepared from 30- to 50-day-old mice (C57BL/6J; The Jackson Laboratory), as previously described (21, 22).The care and use of animals followed all federal and institutional guidelines, the Institutional Animal Care and Use Committees of the Boston Veterans Affairs (VA) Healthcare System, and the Subcommittee on Research Animal Care of the Massachusetts General Hospital. The mice were deeply anesthetized with isoflurane and decapitated. The brains were removed immediately after death, and a section of the brain containing the visual cortex V1 (0.5-1mm anterior from the lambdoid suture) was isolated on ice in a 0° to 5°C oxygenated solution containing 1.25 mM NaH2PO4, 2.5mM KCl, 25mM NaHCO3, 1 mM MgCl2, 25 mM glucose, and 225 mM sucrose, equilibrated with 95% O2–5% CO2 (pH 7.4). This cold solution, with a low sodium ion and without calcium ion content, improved tissue viability. In the same medium, 300- to 400-μm-thick coronal slices were prepared using a vibrating blade microtome (Vibratome 3000 Plus, Ted Pella Inc.) and were incubated at room temperature in an artificial cerebrospinal fluid (aCSF) solution containing 125 mM NaCl, 1.25 mM NaH2PO4, 2.5 mM KCl, 25 mM NaHCO3, 1 mM MgCl2, 2 mM CaCl2, and 25 mM glucose, equilibrated with 95% O2–5% CO2 (pH 7.4). After a 2-hour recovery period, slices that contained V1 were transferred and mounted, caudal side down, to a plastic recording chamber (RC- 27L,Warner Instruments, LLC) with a plastic slice anchor (SHD-27LP/2, Warner Instruments, LLC). The chamber was maintained at 30° ± 2°C and continuously superfused (3.3 ml/min) with oxygenated aCSF solution.

### Patch Clamp Validation

A patch clamp amplifier (MultiClamp 700B, Molecular Devices) operating in voltage-clamp mode with a holding potential of 0 mV was used to measure the artifact signals that were generated during the passage of current through an individual coil for bench testing experiments. The magnitude of the stimulus artifact (pA) was defined as the distance between positive and negative phases of the recorded signal.

### Fluorescence Microscopy

Brain slices were prepared and maintained as described above and were then incubated in a dark room at room temperature in aCSF solution. After a 2-hour recovery period, slices that contained the primary visual cortex (V1) were transferred and mounted, caudal side down, to the plastic recording chamber (RC-27L) with a plastic slice anchor (SHD-27LP/2). Imaging was performed with a Nikon Eclipse FN1 microscope (Nikon Instruments Inc.) through a 20× 0.5 numerical aperture objective (Nikon Fluor 20× /0.50 water immersion objective). The excitation light source (X-Cite 120Q; Excelitas Technologies Corp.) was coupled to the epifluorescent port of the microscope. Calcium fluorescence changes were captured with a sCMOS camera (USB 3.1 Gen 1; 2048 x 2048; 30 frames/s; PCO.Panda 4.2). Images were recorded and then processed using image analysis software (ImageJ; National Institutes of Health). Outlines around the somas of individual pyramidal neurons were defined to create ROIs. The calcium fluorescence transients for individual neurons were calculated as (Δ*F*/*F*_0_) = (*F* − *F*_0_)/*F*_0_, where *F*_0_ was the baseline fluorescence level calculated by averaging over 2s before the onset of stimulation, and subsequently normalized to max observed fluorescence change.

### Data Analysis

MATLAB’s ‘fitnlm’ function was utilized to fit each dose-response curve with a four-parameter sigmoidal function defined by 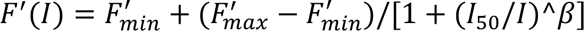, where *F*‘ = (Δ*F*/*F*_0_) and *β* is the dynamic range. *I*_50_is the current amplitude required to elicit 50% of the maximum fluorescence change; this was used as an estimate for the threshold of magnetic stimulation.

## Acknowledgements

This research was supported by NSF CAREER 2239557 (to B.P.T.), AHA 23TPA1057212 (to B.P.T), NIH R21EB034527 (to B.P.T.), BRAIN R01-NS110575 (to S.I.F) and DOD/CDMRP VR170089 (to S.I.F)

